# Serum and Plasma Sphingolipids as Biomarkers of Anthracycline-Induced Cardiotoxicity

**DOI:** 10.1101/2025.01.06.631494

**Authors:** Samia Mohammed, Andreas P. Kalogeropoulos, Victoria Alvarado, Michelle Weisfelner-Bloom, Christopher J. Clarke

## Abstract

Although effective as a chemotherapy, the utility of Doxorubicin (Dox) is hampered by cardiotoxicity. Despite this, the ability to predict and guide monitoring of patients receiving Dox or other anthracyclines is hampered by a lack of effective biomarkers to identify susceptible patients, and to detect early signs of subclinical cardiotoxicity. Based on their well-established roles in the response to Dox and other chemotherapies, we performed a retrospective analysis of serum and plasma sphingolipids (SLs) from patients undergoing anthracycline-containing therapy, correlating with cardiac parameters assessed by echocardiography. Results showed there were substantial changes in both plasma and serum SL species during therapy including ceramide (Cer), deoxydihydroCer, and dihydrosphingosine with reversion towards baseline following treatment. Linear mixed-effects model analysis revealed that at baseline, a number of SLs correlated with adverse cardiac outcomes with serum sphingosine-1-phosphate (S1P) and dihydroS1P, and plasma Cer performing comparably to the prognostic value of pro-NT-BNP, an established biomarker of cardiotoxicity. Intriguingly, while pro-NT-BNP had no predictive value at mid- and post-therapy timepoints, serum S1P and dhS1P and plasma Cer levels showed correlation with adverse outcomes, particularly at the post-therapy timepoint. Finally, analysis of plasma and serum C16:C24- Cer ratios – previously reported as predictive of adverse cardiac outcomes – showed no correlation in the context of anthracycline treatment. Taken together, this pilot study provides supporting evidence that plasma and serum SLs may have benefit as both prognostic and diagnostic biomarkers for patients undergoing anthracycline-containing therapy. This suggests that diagnostic SL measurements – recently implemented for metabolic-associated cardiac disorders – could have wider utility.

---

Although widely used as chemotherapeutics, the clinical utility of Doxorubicin (Dox) and other anthracyclines are limited by dose-dependent cardiotoxicity [1–3]. Although reports vary, dose dependent cardiomyopathy has been reported to range from 7% at 150 mg/m^2^ to up to 65% at 550 mg/m^2^. Dox-induced cardiotoxicity presents as a spectrum of left ventricular dysfunction and clinical heart failure [1–3]. Cardiotoxicity commonly occurs within 12-months of initial treatment but can also present after significant delay [4, 5]. The cardiac damage associated with Dox treatment is considered largely irreversible and can profoundly impact the survival and prognosis of cancer patients [1–3, 6, 7]. Consequently, as cancer survival rates improve and there are increasing number of patients exposed to anthracyclines, strategies that can better predict and control this cardiotoxicity are crucial.

Echocardiography is currently the standard of care for monitoring cardiac function in patients with cancer, owing to widespread availability and safety. The addition of contrast enhanced and three-dimensional echocardiography, have improved subclinical detection of LV dysfunction (LVD) (8). Global longitudinal strain (GLS), a measure of myocardial contractility, provides additional predictive value for LVEF decline in patients receiving anthracyclines (9, 10). As such, European Society of Cardiology Guidelines and recent cardiac imaging guidelines recommend the addition of transthoracic echocardiography (TTE) with GLS as part of the cardiac imaging assessment of patients receiving anthracycline-based chemotherapy (11). Nonetheless, despite improvements in imaging technology, there is still limited data to inform appropriate timing and interval for surveillance imaging prior to, during, and following anthracycline therapy. Additionally, while imaging can detect early signs of damage, there are still limited means for defining patient populations with higher susceptibility or predisposition to anthracycline toxicity. The use of serum or plasma biomarkers holds significant potential, both for the identification of at-risk patient populations as well as for monitoring of cardiotoxicity during treatment. Current biomarkers for anthracycline-induced cardiotoxicity include cardiac troponins (TnI, TnT) and N-terminal pro-BNP peptide (NT-BNP) with increased levels predictive of future LVD (12-17). Some studies have also suggested these biomarkers can have negative predictive value with the absence of changes in their levels reflecting minimal changes in cardiac function (16, 17). Despite this, both biomarkers have limited and ill-defined roles in the prediction of subclinical cardiotoxicity and significant changes in the levels of each biomarker may only be detected after cardiac damage has already occurred (12-17). Consequently, novel biomarkers might add additional value in risk prediction.

Sphingolipids (SL) such as ceramide (Cer), sphingosine (Sph), and Sph-1-phosphate (S1P) are a family of bioactive lipids implicated in many cellular processes (18, 19). Cellular SL levels are controlled by an interlinked network of metabolic enzymes, and alterations in SL metabolism are widely associated with the cellular response to numerous chemotherapies across diverse cancers (20-26). Pathologically, dysregulation of SL metabolism and signaling is linked with a host of diseases including cancers, neurological diseases, and metabolic syndromes with some reports showing that levels of SLs and SL enzymes can correlate with disease prognosis (18, 19, 27-29). In addition to their cellular functions, many SL species are readily detected in serum and plasma. In this context, S1P is very well studied, complexing with HDL and ApoM and being able to influence signaling and functions of endothelial cells through binding to extracellular S1P receptors (30, 31). There is also growing evidence that variations in serum and plasma SL levels are associated with pathological conditions including atherosclerosis (32), sepsis (33), metabolic syndrome (34), and age-related macular degeneration (35). More recently, serum SL profiles were reported to predict asymptomatic vs. symptomatic COVID-19 (36) while an independent study found that serum SLs were prognostic for COVID disease severity (37). In the context of heart disease and function, studies have established specific Cer species as diagnostic indicators of adverse cardiac events. Research by Petersen and colleagues have reported the predictive value of the ratio of C16-Cer to C24-Cer with a high ratio being associated with increased HF risk and altered LVEF (38, 39). Independently, researchers at the Mayo clinic reported that a ‘ceramide score’ obtained through measurements of C16, C18, C24, and C24:1-Cer was a better predictor of future adverse cardiovascular events than monitoring cholesterol levels (40, 41) and led to the implementation of Cer measurements (the CERAM test) as a diagnostic tool. A similar approach by researchers in Utah found that an inclusive score of specific Cer, dhCer, and SM species was a better predictor of coronary artery disease than cholesterol levels (42). However, despite the central role that SLs play in the chemotherapy response, the effects of anthracyclines on serum and plasma SLs or their association with heart function have yet to be fully investigated.

In this study, we performed a retrospective sphingolipidomic analysis of plasma and serum samples from a pilot cohort of patients with breast cancer receiving anthracycline therapy. We analyzed changes in SL levels at baseline, mid, post, and 6-month post treatment, and correlated lipid levels with subclinical changes in heart function detected by TTE and speckle tracking.

## MATERIALS AND METHODS

### Patient samples

This study is a retrospective analysis of plasma and serum samples obtained from a previous prospective study into potential novel biomarkers of anthracycline-induced cardiotoxicity (43). In the prior study, serial blood and echocardiographic assessment was performed on female patients 18-85 years old with a diagnosis of invasive breast cancer without metastases who were planned for anthracycline-inclusive chemotherapy. Patients were assessed at baseline, mid therapy, post therapy and at 6 month follow up. Of the 31 patients screened, serum samples for 20 patients and plasma samples from 18 patients were obtained for analysis. The Institutional Review Board of Stony Brook University approved both the original study (protocol #922042) and the retrospective analysis (protocol #2020-00116) prior to initiation.

### Sphingolipidomic Analysis and Normalization

For lipid analysis, serum and plasma samples were thawed on ice and lipids were extracted as described previously (44, 45). Briefly, 100-200 μl of serum/plasma was diluted in 2ml RPMI medium and 50μl internal standard was added. Sampes were extracted with 2 ml extraction solvent (isopropanol: ethyl acetate 15:85 v/v), vortexed, and centrifuged (5 min, 2500xg). The upper organic phase (approx. 2ml) was transferred to a fresh glass tube. The remaining lower phase was acidified with 100 μl formic acid (98%) and an additional 2 ml of extraction solvent was added. Samples were vortexed and centrifuged (5 min, 2500xg) and the upper organic phase (approx 2ml) was removed and combined with prior extract. The total extract was dried under nitrogen gas and resuspended in 150ul of mobile phase B (1mM ammonium formate, 0.2% formic acid in methanol) and lipids were analyzed by tandem LC/MS/MS at the Stony Brook lipidomics core. Samples were analyzed for ceramide (Cer), dihydroCer (dhCer), sphingosine (Sph), Sph-1-phosphate (S1P), dihydroSph (dhSph), dhSph-1-phosphate (dhS1P), deoxydhCer, and alpha-hydroxyCer (ahCer). Measured lipid levels were normalized to ml of plasma/serum.

### Statistical Analysis

Continuous variables were described as a median (interquartile range) to ensure valid measures of location and dispersion regardless of distribution, and discrete (binary and categorical) variables were described as frequency (percentage). To examine changes over time for the exposures of interest (lipid species concentrations), and the association with outcomes of interest (echocardiographic variables), we employed linear mixed-effects models with patient intercept as the random effect (i.e., each patient had an individual intercept for each variable of interest). Because of the small sample size, we estimated standard errors and confidence intervals for each model using bootstrapping (1000 replications). Changes between time points for each variable of interest were estimated with appropriate contrasts for the marginal means after estimating the corresponding mixed-effects model. We used unadjusted α = 0.05 as the threshold for statistical significance. Analyses were performed with STATA 18.5 (StataCorp LLC, College Station, TX).

## RESULTS

### Anthracycline therapy is associated with alterations in plasma and serum SL levels

Changes in cellular SL levels are a well-established hallmark of the response to many chemotherapies (20-26). In contrast, few studies have examined effects of chemotherapies on SLs in normal tissue, nor in patient samples. Here, we performed a retrospective sphingolipidomic analysis of a cohort of plasma (18 total) and serum samples (20 total) collected at baseline, mid-therapy, end-therapy and at 6-mo follow up from breast cancer patients undergoing anthracycline therapy. Lipid profiles for plasma (**Fig. 1**) and serum (**Fig. 2**) are summarized. Of note, highly similar profiles were observed between serum and plasma for all lipids analyzed with the primary difference being in the absolute levels of the lipids. In this context, plasma samples tended to have lower absolute levels for all lipids with the exceptions of dhCer and dhSph, which were higher in plasma than serum. For Cer, deoxydhCer and dhSph, levels in both plasma and serum increased throughout therapy, peaking at the post-therapy time point and decreasing towards baseline at follow up. For S1P and dhS1P, levels in both plasma and serum tended to decrease with therapy and remain somewhat flat throughout post therapy and follow up. Similar decreases with therapy were observed with both plasma and serum ahCer but at follow up, levels had begun to increase towards baseline. For dhCer, levels remained somewhat flat throughout therapy in both plasma and serum but sharply decreased at 6 mo follow up. Finally, levels of Sph tended to increase with therapy and return towards baseline at follow up but where were differences as to the peak levels – with serum Sph peaking in mid therapy and plasma Sph peaking at follow up. Given the small sample size, the confidence interval for lipid values is broad but there were nonetheless some statistically significant changes observed. In serum, there were significant changes in deoxydhCer at midpoint (p = 0.0034) and post therapy (p = 0.0045) vs. baseline levels, Cer at post therapy vs. baseline (p = 0.0005), and dhSph at post therapy vs. baseline (p = 0.016). In plasma, there were similar significant changes in deoxydhCer at midpoint (p = 0.0087) and post therapy (p = 0.0008) vs. baseline levels, Cer at post therapy vs. baseline (p = 0.0099), and dhSph at post therapy vs. baseline (p = 0.0202). There were also significant changes in dhS1P at mid therapy vs baseline (p = 0.0142) and Sph at post-therapy vs baseline (p = 0.0229). Taken together, these results show that alterations in serum and plasma SL levels are associated with anthracycline-containing therapy regimens.

**Figure 1.**
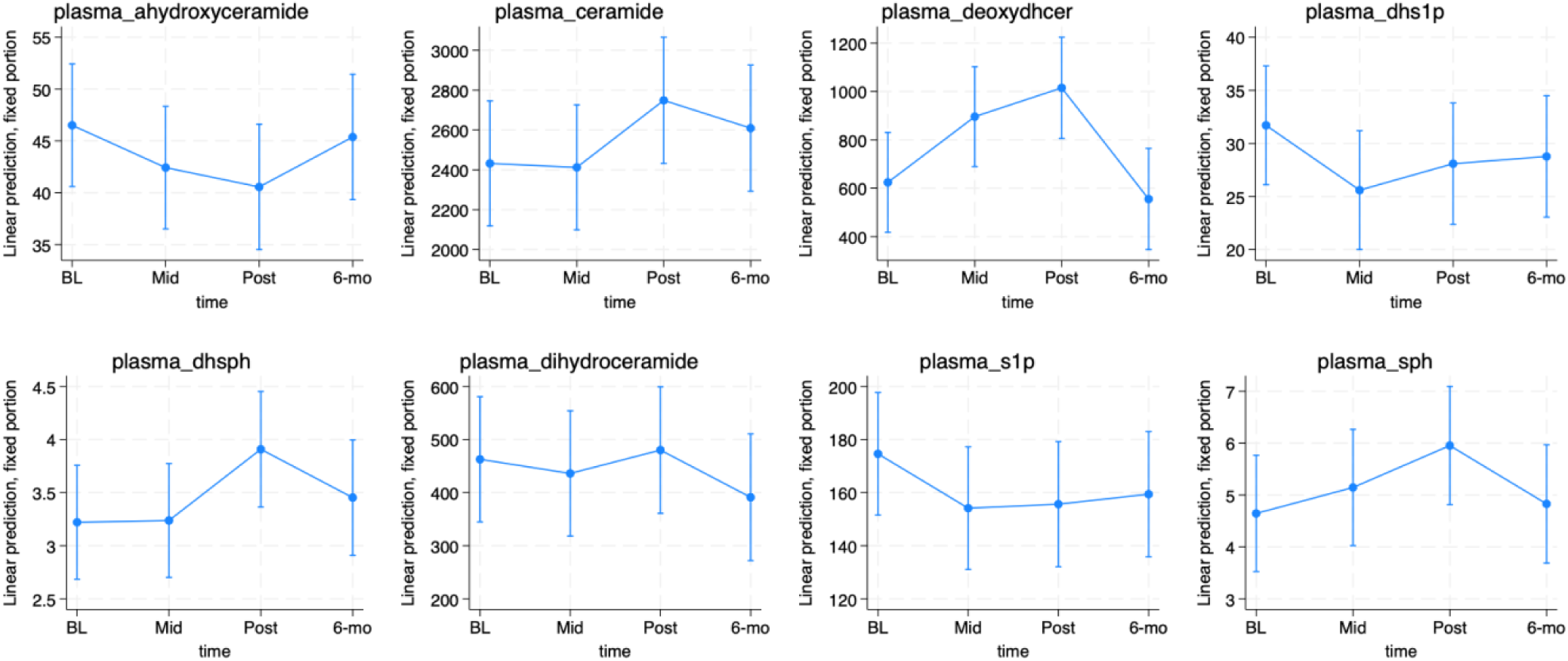
Alterations in plasma sphingolipid levels in patients undergoing anthracycline-containing therapy: Lipids were extracted from 100-200ul of plasma and analyzed for the sphingolipid classes shown as described in “Materials and Methods”. Data were normalized to ml plasma and pmol/ml at baseline (BL), midpoint (Mid), Post-therapy (post) and 6-month follow up (6-mo). Data are expressed as mean +/- 95% confidence interval.

**Figure 2.**
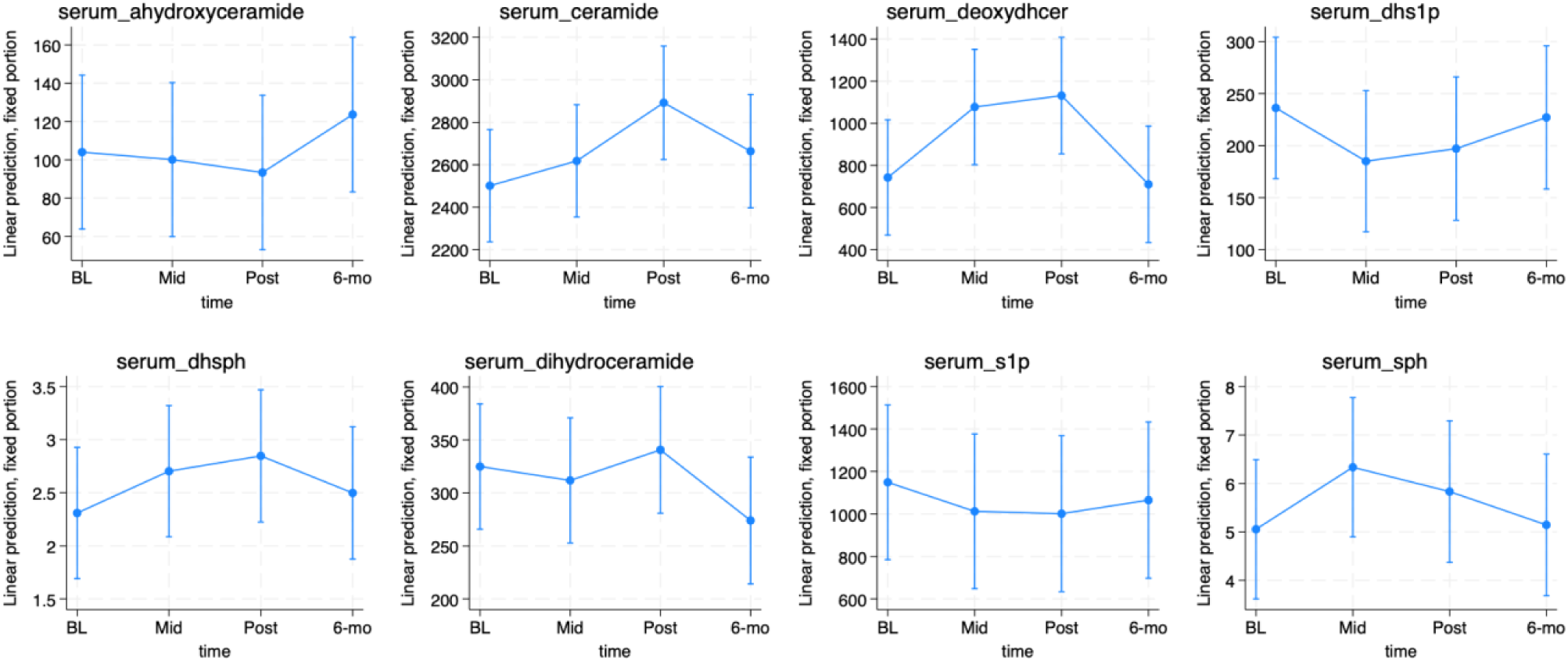
Alterations in serum sphingolipid levels in patients undergoing anthracycline-containing therapy: Lipids were extracted from 100-200ul of serum and analyzed for the sphingolipid classes shown as described in “Materials and Methods”. Data were normalized to ml plasma and pmol/ml at baseline (BL), midpoint (Mid), Post-therapy (post) and 6-month follow up (6-mo). Data are expressed as mean +/- 95% confidence interval.

**Figure 3.**
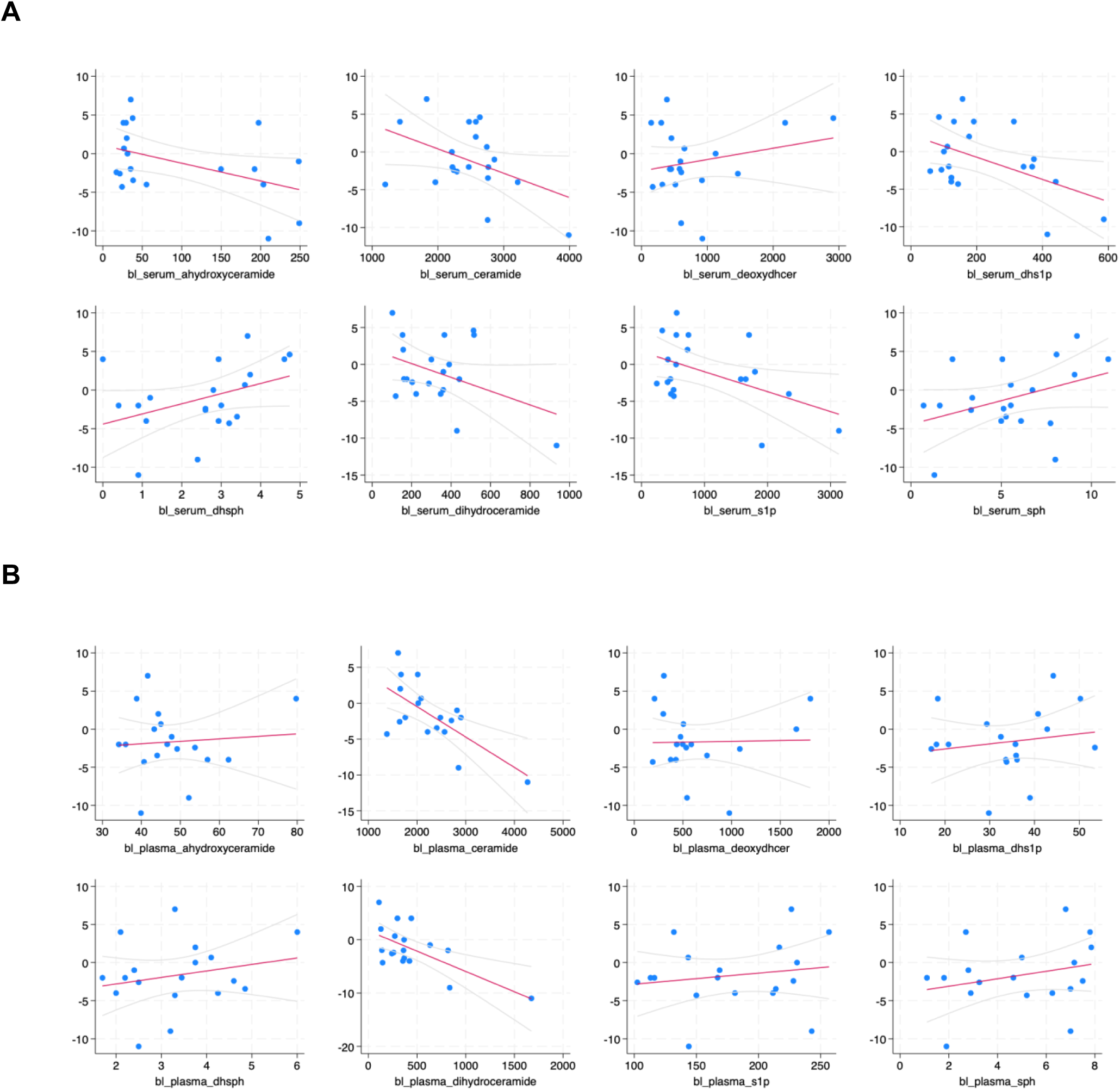
Baseline plasma and serum sphingolipid levels correlated with changes in LVEF in patients receiving anthracycline-containing therapy: Linear mixed-effects models were used to correlate baseline lipid levels for (A) serum and (B) plasma with changes in LVEF from baseline to 6 mo follow up. Here, a decrease in LVEF would be considered an adverse outcome. Trendline is shown in red with 95% confidence interval shown in light gray.

**Figure 4.**
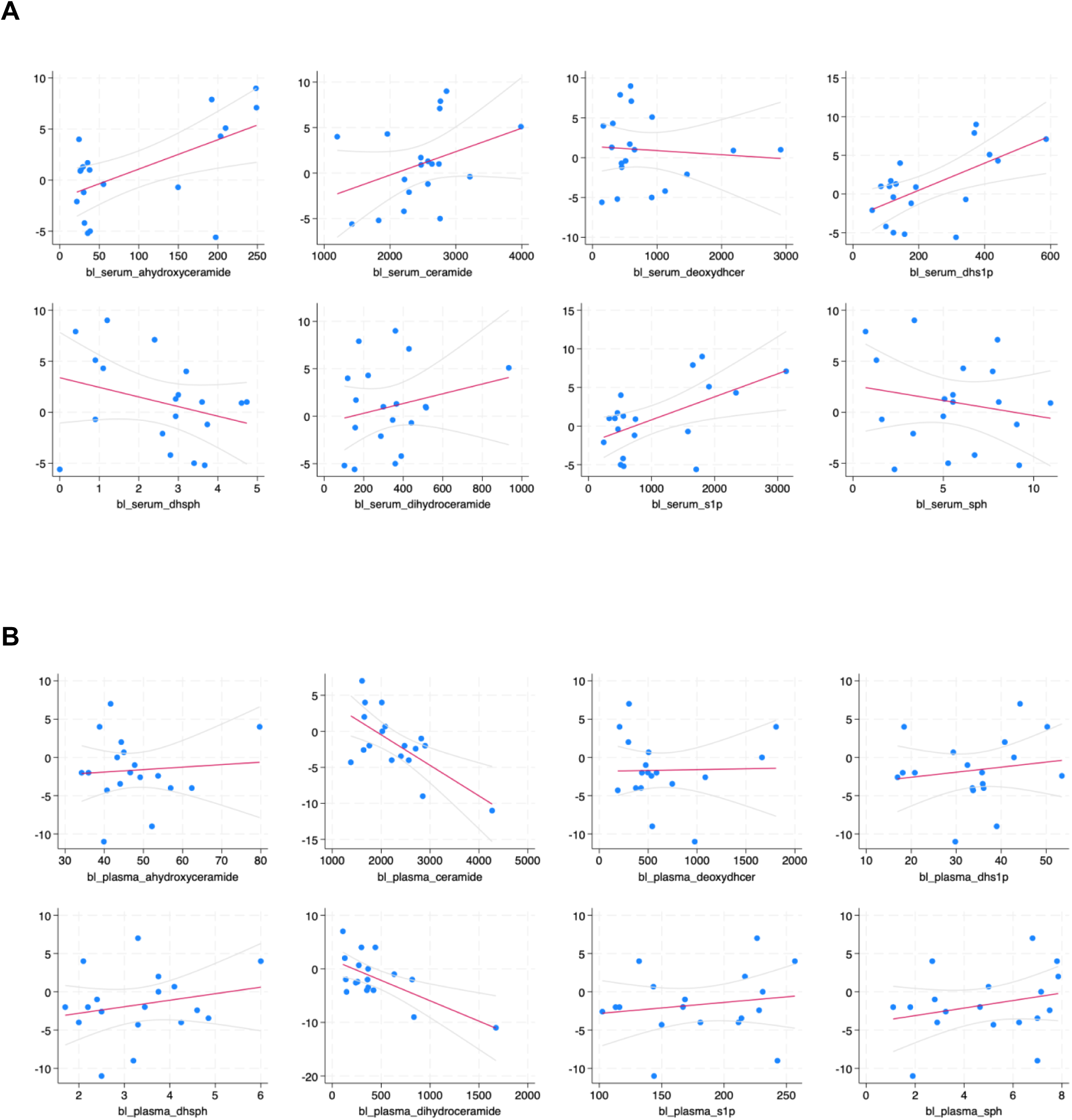
Baseline plasma and serum sphingolipid levels correlated with changes in GLS in patients receiving anthracycline-containing therapy: Linear mixed-effects models were used to correlate baseline lipid levels for (A) serum and (B) plasma with changes in GLS from baseline to 6 mo follow up. Here, am increase in GLS would be considered an adverse outcome. Trendline is shown in red with 95% confidence interval shown in light gray.

### Correlation of baseline SL levels with adverse changes in cardiac function (LVEF, GLS)

Prior studies have associated circulating levels of SLs (mainly Cer) with increased likelihood of HF and adverse cardiovascular outcomes (36-40). Accordingly, we reasoned that baseline SL levels may be associated with an increased susceptibility to Dox-induced cardiac damage. To assess this, we asked if baseline SL levels in serum or plasma were associated with subsequent average changes in LVEF and GLS as measured by echocardiography and speckle tracking at the same time points. Here, it is important to note that adverse cardiac function is reflected by a decrease in LVEF (e.g. from 70% to 60%) and/or an increase in GLS (e.g. -20% to -15%). Consequently, if elevated levels of a lipid are associated with adverse heart function, then we would expect a negative correlation with LVEF and a positive correlation with GLS. With LVEF, there were significant negative correlations with baseline serum levels of S1P (r = -0.485, p = 0.038) and dihydroS1P (r = -0.489, p = 0.034) with a negative, but marginally insignificant correlation observed for ahCer (r = -0.449, p = 0.067). In contrast, there were negative correlations of LVEF with baseline plasma levels of Cer (r = -0.671, p = 0.002) and dhCer (r = -0.652, p = 0.008). Analysis of baseline lipid levels with changes in GLS found significant correlations with serum S1P (r = 0.555, p = 0.007) and dihydroS1P (r = 0.601, p = 0.001) consistent with LVEF results. Notably, here there were also significant correlations with ahCer (r = 0.580, p = 0.011). For baseline plasma levels, there were positive correlations of GLS changes with Cer (r = 0.537, p = 0.046), consistent with LVEF correlations, but there was no significant correlation of GLS with dhCer levels (r = 0.415, p = 0.168). To provide a point of comparison, we correlated levels of pro-NT-BNP – an established biomarker of anthracycline cardiotoxicity that was previously investigated in this cohort (43) – with subsequent changes in LVEF and GLS from baseline to post therapy and 6 month follow up. This analysis revealed that baseline pro-NT-BNP levels negatively correlated with LVEF changes but this was not quite statistically significant (r = -0.469, p = 0.082). However, there was a significant correlation between pro-NT-BNP and GLS changes (r = 0.596, p = 0.001) (**Table 1**). Taken together, these results suggest that both plasma and serum SL levels can be predictive of anthracycline-driven changes in LVEF and GLS, and that they function comparably to an existing biomarker.

**Table 1:** Correlation of the established biomarker pro-NT-BNP with cardiac parameters at baseline, mid-therapy, and post-therapy timepoints. Shown are corresponding beta value and p-value for correlation of pro-NT-BNP at the respective timepoints with changes in LVEF and GLS as shown.

|  | LVEF |  | GLS |  |
| --- | --- | --- | --- | --- |
|  | Beta | p-value | Beta | p-value |
| Baseline | -0.469 | 0.082 | 0.596 | 0.001 |
| Mid-therapy | 0.014 | 0.956 | 0.156 | 0.510 |
| Post | -0.127 | 0.660 | 0.282 | 0.267 |

### Correlation of mid and post-therapy SL levels with adverse changes in cardiac function (LVEF, GLS)

Prognostic biomarkers can be informative for identifying those patients who may have increased likelihood of anthracycline-induced cardiotoxicity. However, an early indicator of subclinical cardiotoxicity would also have value as this could provide guidance for monitoring of cardiac function, administering of drugs that can try to preserve heart function, or dose reduction of anthracyclines. For similar reasons, a post-therapy biomarker that is predictive of subsequent adverse cardiac changes would also have value. Notably, in this patient cohort, while pro-NT-BNP levels had been previously observed to increase during treatment (43), mid therapy levels of pro- NT-BNP were not predictive of subsequent changes for LVEF (r = 0.014, p = 0.956) or GLS (r = 0.156, p = 0.510). Similarly, post-therapy levels of pro-NT-BNP did not correlate well with subsequent changes in LVEF (r = -0.127, p = 0.660) or GLS (r = 0.282, p = 0.267) although results were marginally better for GLS than for LVEF. As correlation of baseline SL levels with adverse LVEF and GLS outcomes showed promise and changes in both plasma and serum SL levels during therapy were observed, we hypothesized that midpoint and post therapy SLs might also have predictive value for LVEF and GLS changes.

Analysis of serum SL levels with LVEF revealed no significant correlations with midpoint lipid values – although ahCer (r = -0.414, p = 0.064) was borderline. However, at the post-chemotherapy endpoint, there were significant negative correlations with serum S1P levels and LVEF (r = -0.488, p = 0.016) with ahCer again showing borderline significant (r = -0.425, p = 0.054). In contrast, comparisons of serum SL levels with GLS showed more promising results; at midpoint, there were significant correlations of S1P (r = 0.570, p = 0.005), dihydroS1P (r = 0.569, p = 0.007) and ahCer (r = 0.583, p = 0.020) although at the post-chemotherapy time-point, there was only significant correlations with S1P (r = 0.500, p = 0.034) and, again, borderline significance for ahCer (r = 0.486, p = 0.053). For plasma SL levels, comparisons with LVEF revealed no significant negative correlations with midpoint lipid values – although intriguingly there were positive correlations (although not quite statistically significant) for dhSph (r = 0.478, p = 0.083) and dhS1P (r = 0.444, p = 0.088). However, at the post-chemotherapy endpoint, there was a significant negative correlation of LVEF with plasma Cer (r = -0.527, p = 0.019). Comparison of plasma SL levels with GLS also showed no positive correlations at midpoint although surprisingly, there were significant negative correlations with Sph (r = -0.448, p = 0.043) and dhSph (r = -0.550, p = 0.005). However, at endpoint, there was a significant positive correlation with Cer (r = 0.516, p = 0.047) and a negative correlation with dhS1P (r = -0.483, p = 0.041). Taken together, these results suggest that plasma and serum SLs can have prognostic value for adverse cardiac outcomes both during and immediately post therapy – although results were more robust with GLS than LVEF. They also suggest that some SLs could also be positive indicators of cardiac function during anthracycline therapy.

### The C16:C24-Cer ratio does not correlate with adverse changes in cardiac function in anthracycline patients

Prior research has reported that the ratio of plasma C16-Cer to C24-Cer is a predictor of adverse cardiac outcomes superior to cholesterol levels (38), with follow-up studies suggesting that a higher C16: C24 ratio was associated with lower LVEF and lower global circumferential strain but was not associated with changes in GLS (39). Given the results above, we were curious if the C16: C24-Cer ratio may be similarly predictive of Dox-induced cardiotoxicity and assessed correlation of serum and plasma C16:C24-Cer at baseline, mid therapy, and post therapy with LVEF and GLS as above (**Table 2**). Analysis of the serum C16: C24-Cer ratio showed no significant correlation with changes in LVEF at baseline (r = -0.099, p = 0.798), midpoint (r = -0.271, p = 0.329), or post chemotherapy (r = -0.436, p = 0.098) although it should be noted that the correlation was approaching significance at post-therapy. Comparison of serum C16: C24-Cer ratio with GLS showed similar trends with no significant correlations at baseline (r = 0.013, p = 0.978), midpoint (r = 0.275, p = 0.364), or post chemotherapy (r = 0.287, p = 0.268) although in contrast with LVEF, there was less improvement at later timepoints. Analysis of plasma C16:C24-Cer ratios showed there were no significant correlations for changes in LVEF at baseline (r = 0.104, p = 0.719), midpoint (r = -0.175, p = 0.466), or post chemotherapy (r = 0.052, p = 0.816). Similar results were obtained for changes in GLS at baseline (r = -0.046, p = 0.861), midpoint (r = 0.151, p = 0.526), or post chemotherapy (r = -0.172, p = 0.485). Taken together, these results suggest that the C16:C24-Cer ratio – either in serum or plasma – is not predictive of cardiotoxic effects of anthracycline therapy.

**Table 2:** Correlation of serum sphingolipid levels with cardiac parameters at mid-therapy and post-therapy timepoints. Shown are corresponding beta value and p-value for correlation of serum sphingolipid levels at mid- and post-therapy timepoints with subsequent changes in LVEF and GLS.

|  | Mid LVEF |  | Mid GLS |  | Post LVEF |  | Post GLS |  |
| --- | --- | --- | --- | --- | --- | --- | --- | --- |
|  | Beta | p-value | Beta | p-value | Beta | p-value | Beta | p-value |
| Cer | -0.160 | 0.368 | 0.131 | 0.597 | -0.330 | 0.177 | 0.254 | 0.238 |
| dhCer | 0.095 | 0.679 | 0.164 | 0.463 | -0.044 | 0.886 | 0.105 | 0.632 |
| S1P | -0.317 | 0.149 | 0.570 | 0.005 | -0.488 | 0.016 | 0.500 | 0.034 |
| dhS1P | -0.190 | 0.274 | 0.569 | 0.007 | -0.438 | 0.100 | 0.439 | 0.177 |
| Sph | 0.273 | 0.302 | -0.246 | 0.272 | -0.191 | 0.535 | 0.094 | 0.741 |
| dhSph | 0.372 | 0.134 | -0.341 | 0.124 | 0.044 | 0.872 | -0.124 | 0.665 |
| deoxydhCer | 0.283 | 0.245 | -0.138 | 0.514 | 0.229 | 0.348 | -0.077 | 0.732 |
| ahCer | -0.414 | 0.064 | 0.583 | 0.020 | -0.425 | 0.054 | 0.486 | 0.053 |

**Table 3:** Correlation of plasma sphingolipid levels with cardiac parameters at mid-therapy and post-therapy timepoints. Shown are corresponding beta value and p-value for correlation of plasma sphingolipid levels at mid- and post-therapy timepoints with subsequent changes in LVEF and GLS.

|  | Mid LVEF |  | Mid GLS |  | Post LVEF |  | Post GLS |  |
| --- | --- | --- | --- | --- | --- | --- | --- | --- |
|  | Beta | p-value | Beta | p-value | Beta | p-value | Beta | p-value |
| Cer | -0.181 | 0.392 | 0.339 | 0.220 | -0.527 | 0.019 | 0.516 | 0.047 |
| dhCer | -0.163 | 0.388 | 0.427 | 0.110 | -0.367 | 0.152 | 0.348 | 0.133 |
| S1P | 0.243 | 0.333 | 0.006 | 0.986 | 0.214 | 0.439 | -0.367 | 0.151 |
| dhS1P | 0.444 | 0.088 | -0.165 | 0.541 | 0.384 | 0.142 | -0.483 | 0.041 |
| Sph | 0.457 | 0.153 | -0.448 | 0.043 | 0.075 | 0.817 | -0.303 | 0.356 |
| dhSph | 0.478 | 0.083 | -0.550 | 0.005 | 0.086 | 0.763 | -0.317 | 0.331 |
| deoxydhCer | 0.240 | 0.380 | -0.152 | 0.533 | 0.190 | 0.549 | -0.060 | 0.841 |
| ahCer | -0.181 | 0.512 | 0.192 | 0.488 | -0.097 | 0.687 | 0.090 | 0.779 |

**Table 4:**
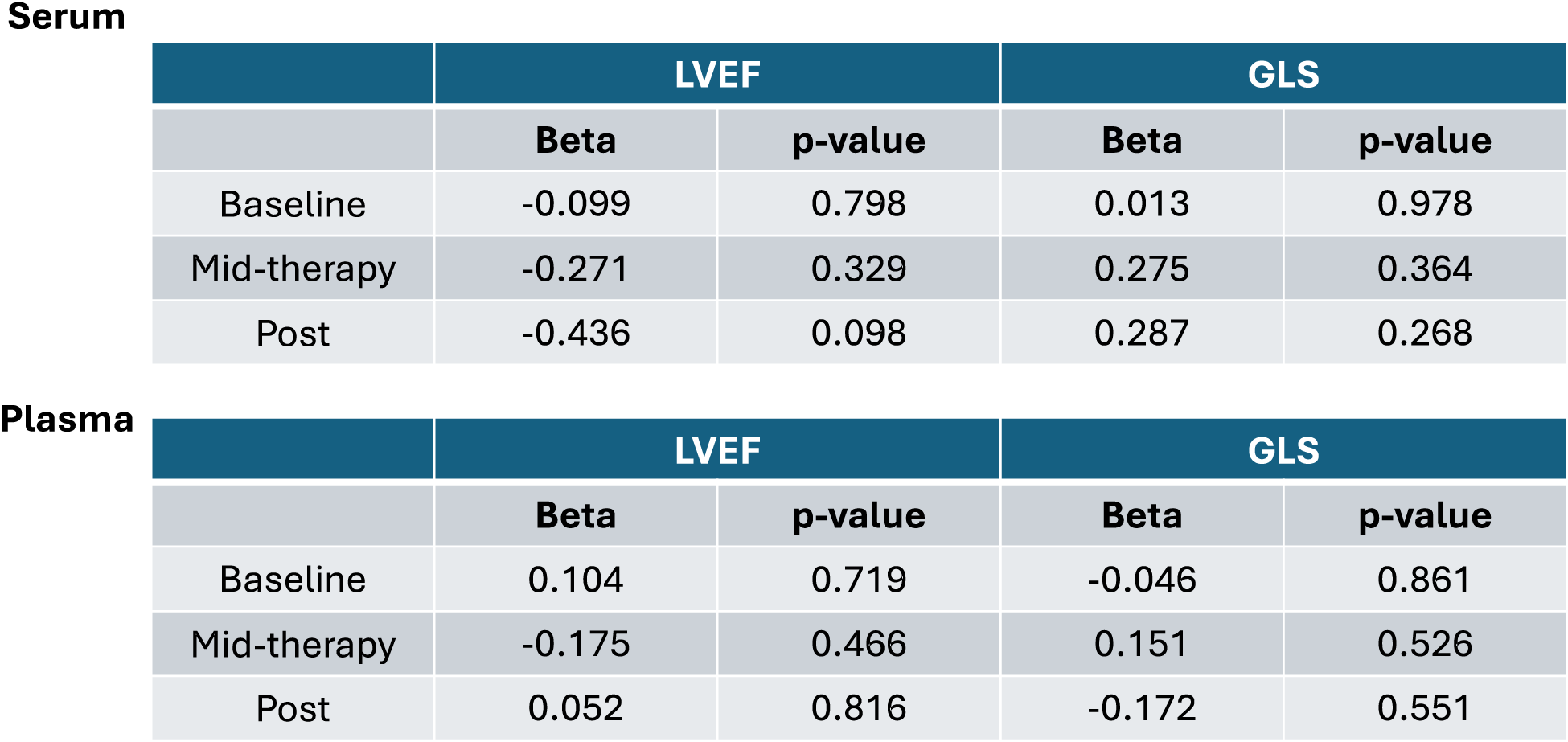
Correlation of serum and plasma C16:C24-Cer ratio with cardiac parameters at baseline, mid-therapy and post-therapy timepoints. Shown are corresponding beta value and p-value for correlation of serum amd plasma C16:C24-Cer ration at the time points shown with subsequent changes in LVEF and GLS.

**Serum**
|  | LVEF |  | GLS |  |
| --- | --- | --- | --- | --- |
|  | Beta | p-value | Beta | p-value |
| Baseline | -0.099 | 0.798 | 0.013 | 0.978 |
| Mid-therapy | -0.271 | 0.329 | 0.275 | 0.364 |
| Post | -0.436 | 0.098 | 0.287 | 0.268 |

**TABLE 4**
|  | LVEF |  | GLS |  |
| --- | --- | --- | --- | --- |
|  | Beta | p-value | Beta | p-value |
| Baseline | 0.104 | 0.719 | -0.046 | 0.861 |
| Mid-therapy | -0.175 | 0.466 | 0.151 | 0.526 |
| Post | 0.052 | 0.816 | -0.172 | 0.551 |

## DISCUSSION

In this study, we performed a retrospective analysis of SL levels in serum and plasma samples from anthracycline-treated patients, correlating lipid levels with LVEF and GLS parameters of cardiac function. Our results show robust changes in plasma and serum SLs during anthracycline-containing regimens, with Cer, deoxydhCer, and dhSph being the most notable species affected. Furthermore, we found correlations of plasma and serum SLs at baseline, mid-therapy, and post-therapy with adverse changes in cardiac function. In contrast, the ratio of C16:C24-Cer, previously found to be a predictor of worse cardiac outcomes, was not predictive of anthracycline-induced effects. Taken together, this pilot study offers preliminary evidence in support of plasma and serum SLs as potential biomarkers of interest for anthracycline-induced cardiotoxicity.

Since the discovery of their signaling functions, the biological roles of SLs have been the subject of intensive research and alterations in cellular SLs are now well-established components of the response to various chemotherapies (20-26). While this has led to interest in modulating SLs to enhance therapeutic efficacy in cancer cells, comparatively few studies have looked at the effects of chemotherapies on SLs in non-transformed tissues, nor in plasma or serum of chemotherapy-treated patients. Here, our results show that alterations in plasma and serum SL levels occur during anthracycline-containing therapy regimens. These changes were broad, occurring across multiple SL species, and despite being from a small cohort, some of these changes reached statistical significance – particularly for deoxydhCer, Cer, and dhSph in both plasma and serum. Lipid levels were also clearly driven by treatment as levels increased/decreased at mid therapy and post therapy time points before reverting towards the baseline levels at 6-month of follow up. Thus, as with many other pathological conditions (32-37), chemotherapy treatment can drive systemic changes in plasma and serum SLs. Although levels were used to correlate with cardiac function, the source of these lipids is unclear as anthracyclines can also affect vascular endothelial cells (46, 47) and other tissues (48, 49) as well as the cancer cells themselves (21, 22). Moreover, these changes cannot be attributed solely to anthracyclines, particularly as SL metabolism can be affected by taxanes (25, 26), which are used in combination with anthracyclines for BC treatment.

While echocardiography remains the standard of care for cardiac monitoring, serum and plasma biomarkers hold significant potential for identifying at-risk populations prior treatment as well as in the detection of subclinical toxicity during chemotherapy. Here, our results show that both plasma and serum levels of a number of SL species were correlated with subsequent decreases in cardiac function as indicated by decreased LVEF and GLS. Notably, correlations of baseline SL species with adverse outcomes were at least comparable to or better than the existing biomarker pro-NT-BNP at predicting adverse changes in GLS and LVEF (and NT-BNP was not quite statistically significant for the latter). However, at mid-therapy and post-therapy timepoints, the levels of SL species – primarily S1P, dhS1P, and ahCer in serum, and Cer in plasma – were better at predicting subsequent adverse GLS or LVEF changes; indeed, mid and post-therapy pro-NT-BNP levels in this cohort were not predictive for either parameter. While baseline correlations can provide guidance on patient monitoring, correlations during treatment can be equally valuable for indicating whether cardioprotective intervention should be initiated, particularly as studies have shown that the earlier the intervention, the more substantial the recovery of LV function (50, 51). As there is evidence that changes in GLS can be more indicative of early subclinical cardiotoxicity (9, 10), these correlations may be of more clinical relevance. In this context, the correlations of SL levels with adverse GLS changes were more robust than those with LVEF – particularly at the mid-therapy time-point. An additional point of interest here is that midpoint plasma levels of Sph and dhSph showed a negative correlation with GLS (Sph – r = -0.448; dhSph –r = -0.550) and positive correlations with LVEF (albeit not statistically significant). Similar correlations for GLS were also observed at post-therapy with plasma dhS1P levels (GLS - r = -0.483). As these correlations would be more indicative of maintained or improved cardiac function, this raises the intriguing possibility that serum or plasma SLs could function both as negative and positive indicators of cardiac function during therapy.

While our focus in this study was on total lipid levels, there is the possibility that individual lipid species from each family may have comparable or better predictive power and serve to drive the broader correlations seen with total lipids. Indeed, the Cer score from Mayo clinic centers on C16, C18, C24, and C24:1-Cer species (40, 41). However, given the number of potential SL species that could be measured (both in serum and plasma), we considered this more complex analysis as beyond the scope of this initial study. Finally, the lack of correlation of the C16:C24-Cer ratio with adverse outcomes suggests that the chemotherapy effects on the heart might be distinct from those associated with more general adverse cardiac health changes. Nonetheless, we do note that at post-chemotherapy time points, the serum C16: C24-Cer ratio showed a negative correlation with subsequent LVEF changes and was approaching significance. Thus, it is plausible that analysis of a larger patient cohort may bring these results in line with prior reports. Along these lines, although these results show promise for SLs as biomarkers, there are limitations of this study that are important to note. First and foremost, this was a pilot study and limited to 18 sets of plasma and 20 sets of serum; thus, the study does not have sufficient power to detect major changes in biomarkers, SL levels, or cardiac changes. An important aspect related to this is that the vast majority (c. 90%) of the obtained samples were from a Caucasian population and as well as being from patients with few cardiac risk factors; thus, the relevance of these findings to broader populations or to patients who may have an elevated susceptibility to cardiotoxicity is limited and unclear. Finally, samples and cardiac function data were limited to a six-month follow-up period – and given that anthracycline-induced cardiotoxicity commonly occurs up to 12 months post therapy (4, 5), a follow up at later time points could have provided further insight. Overall, a follow up retrospective or future prospective analysis with a larger sample population and keeping these considerations in mind is crucial to extend and further validate these findings.

In conclusion, this study has shown that anthracycline-induced cardiotoxicity regimens are associated with alterations in plasma and serum SLs and provides evidence that plasma and serum SLs have potential to serve as biomarkers for anthracycline-induced cardiotoxicity. Importantly, SL levels were correlated with adverse changes in both GLS and LVEF, performed at least comparably to the existing biomarker pro-NT-BNP, and showed promise both for pre-treatment identification and mid/post therapy detection of patients with increased likelihood of adverse cardiac outcomes. Although preliminary in nature, these results suggest that the use of clinical SL measurements as a diagnostic tool for cardiac function could be applied to patients undergoing treatment with anthracyclines or other known cardiotoxic chemotherapies and may be beneficial for guiding the monitoring and management of cardiac function both during and after therapy.

## ACKNOWLEDGEMENTS

The authors wish to acknowledge the Stony Brook Cancer Center Lipidomics Shared Resource for expert assistance with sphingolipid analysis. We are also extremely grateful to all of the patients who participated in the original study (43), giving both their time and their samples. This work was supported by a Stony Brook Department of Medicine Pilot Project and National Institutes of Health R01 CA248080. The original study (43) was sponsored by Gilead Therapeutics.

## REFERENCES

1. Sawyer DB. Anthracyclines and heart failure. New Engl J Med. 368(12): 1154–1156, 2013. PMID: 23514294.

2. Mitry MA, Edwards JG. Doxorubicin-induced heart failure: phenotype and molecular mechanisms. Int. J. Cardiol Heart Vasc. 10: 17–24, 2016. PMCID: PMC4871279.

3. Ferrans VJ, Clark JR, Zhang J, Yu ZX, Herman EH. Pathogenesis and prevention of doxorubicin cardiomyopathy. Tsitologiia 39: 928–937, 1997. PMID: 9505340.

4. Franco VI, Lipshultz SE. Cardiac complications in childhood cancer survivors treated with anthracyclines. Cardiol Young. 25(Suppl2) 107–116, 2015. PMID: 26377717.

5. Boyd A, Stoodley P, Richards D, Hui R, Harnett P, Vo K, Marwick T, Thomas L. Anthracyclines induce early changes in left ventricular systolic and diastolic function: a single centre study. PLoS One. 12(4): e0175544, 2017. PMCID: PMC5391073.

6. Bloom MW, Vo JB, Rodgers JE, Ferrari AM, Nohria A, Deswal A, Cheng RK, Kittleson MM, Upshaw JN, Palaskas N, Blaes A, Brown SA, Ky B, Lenihan D, Maurer MS, Fadol A, Skurka K, Cambareri C, Chauhan C, Barac A. Cardio-Oncology and Heart Failure: a Scientific Statement From the Heart Failure Society of America. J Card Fail. S1071–9164, 2024.

7. Kero AE, Järvelä LS, Arola M, Malila N, Madanat-Harjuoja LM, Matomäki J, Lähteenmäki PM. Cardiovascular morbidity in long-term survivors of early-onset cancer: a population-based study. Int J Cancer. 134:664–73, 2014. PMID: 23852751.

8. Baldassarre LA, Ganatra S, Lopez-Mattei J, Yang EH, Zaha VG, Wong TC, Ayoub C, DeCara JM, Dent S, Deswal A, Ghosh AK, Henry M, Khemka A, Leja M, Rudski L, Villarraga HR, Liu JE, Barac A, Scherrer-Crosbie M; ACC Cardio-Oncology and the ACC Imaging Councils. Advances in Multimodality Imaging in Cardio-Oncology: JACC State-of-the-Art Review. J Am Coll Cardiol. 80(16): 1560–1578.

9. Oikonomou EK, Kokkinidis DG, Kampaktsis PN, Amir EA, Marwick TH, Gupta D, Thavendiranathan P. Assessment of Prognostic Value of Left Ventricular Global Longitudinal Strain for Early Prediction of Chemotherapy-Induced Cardiotoxicity: A Systematic Review and Meta-analysis. JAMA Cardiol. 4(10): 1007–1018, 2019. PMCID: PMC6705141.

10. Chimed S, Stassen J, Galloo X, Meucci MC, Knuuti J, Delgado V, van der Bijl P, Ajmone Marsan N, Bax JJ. Prognostic Relevance of Left Ventricular Global Longitudinal Strain in Patients With Heart Failure and Reduced Ejection Fraction. Am J Cardiol. 202: 30–40, 2023. PMID: 37413704

11. Lyon AR, López-Fernández T, Couch LS, Asteggiano R, Aznar MC, Bergler-Klein J, Boriani G, Cardinale D, Cordoba R, Cosyns B, Cutter DJ, de Azambuja E, de Boer RA, Dent SF, Farmakis D, Gevaert SA, Gorog DA, Herrmann J, Lenihan D, Moslehi J, Moura B, Salinger SS, Stephens R, Suter TM, Szmit S, Tamargo J, Thavendiranathan P, Tocchetti CG, van der Meer P, van der Pal HJH; ESC Scientific Document Group. 2022 ESC Guidelines on cardio-oncology developed in collaboration with the European Hematology Association (EHA), the European Society for Therapeutic Radiology and Oncology (ESTRO) and the International Cardio-Oncology Society (IC-OS). Eur J Heart Cardiovasc Imaging. 23(10): e333-e465, 2022. PMID: 36017575

12. Herman EH, Lipshultz SE, Rifai N, Zhang J, Papoian T, Yu ZX, Takeda K, Ferrans VJ. Use of cardiac troponin T levels as an indicator of doxorubicin-induced cardiotoxicity. Cancer Res. 58(2): 195–197, 1998. PMID: 9443390

13. Tzolos E, Adamson PD, Hall PS, Macpherson IR, Oikonomidou O, MacLean M, Lewis SC, McVicars H, Newby DE, Mills NL, Lang NN, Henriksen PA. Dynamic Changes in High-Sensitivity Cardiac Troponin I in Response to Anthracycline-Based Chemotherapy. Clin Oncol (R Coll Radiol). 32(5): 292–297, 2020. PMCID: PMC7139216.

14. Suzuki T, Hayashi D, Yamazaki T, Mizuno T, Kanda Y, Komuro I, Kurabayashi M, Yamaoki K, Mitani K, Hirai H, Nagai R, Yazaki Y. Elevated B-type natriuretic peptide levels after anthracycline administration. Am. Heart J. 136, 362–363, 1998. PMID: 9704703

15. Muckiene G, Vaitiekus D, Zaliaduonyte D, Zabiela V, Verseckaite-Costa R, Vaiciuliene D, Juozaityte E, Jurkevicius R. Prognostic Impact of Global Longitudinal Strain and NT-proBNP on Early Development of Cardiotoxicity in Breast Cancer Patients Treated with Anthracycline-Based Chemotherapy. Medicina (Kaunas). 59(5): 953, 2023. PMID: 37241185

16. Michel L, Mincu RI, Mahabadi AA, Settelmeier S, Al-Rashid F, Rassaf T, Totzeck M. Troponins and brain natriuretic peptides for the prediction of cardiotoxicity in cancer patients: a meta-analysis. Eur J Heart Fail 22(2): 350–361, 2020. PMID: 31721381.

17. Bisoc A, Ciurescu D, Rădoi M, Tântu MM, Rogozea L, Sweidan AJ, Bota DA. Elevations in High-Sensitive Cardiac Troponin T and N-Terminal Prohormone Brain Natriuretic Peptide Levels in the Serum Can Predict the Development of Anthracycline-Induced Cardiomyopathy. Am J Ther. 27(2): e142–e150, 2020. PMID: 30648987

18. Hannun YA, Obeid LM. Sphingolipids and their metabolism in physiology and disease. Nat Rev Mol Cell Biol. 19(3): 175–192, 2017. PMID: 29165427

19. Ogretmen B. Sphingolipid metabolism in cancer signaling and therapy. Nat Rev Cancer. 18(1): 33–50, 2018. PMCID: PMC5818153.

20. Snider JM, Trayssac M, Clarke CJ, Schwartz N, Snider AJ, Obeid LM, Luberto C, Hannun YA. Multiple actions of doxorubicin on the sphingolipid network revealed by flux analysis. J Lipid Res. 69(4): 819–831, 2019. PMCID: PMC6446699

21. Liu YY, Yu JY, Yin D, Patwardhan GA, Gupta V, Hirabayshi Y, Holleran WM, Giuliano AE, Jazwinski SM, Gouaze-Andersson V, Consoli DP, Cabot MC. A role for ceramide in driving cancer cell resistance to doxorubicin. FASEB J 22(7): 2541–2551 (2008). PMCID: PMC2883648.

22. Guillermet-Guibert J, Davenne L, Pchejetski D, Saint-Laurent N, Brizuela L, Guilbeau-Frugier C, Delisle MB, Cuvillier O, Susini C, Bousquet C. Targeting the sphingolipid metabolism to defeat pancreatic cancer cell resistance to the chemotherapeutic gemcitabine drug. Mol Cancer Ther. 8(4): 809–20, 2009. PMID:19372554

23. Senkal CE, Ponnusamy S, Rossi MJ, Bialewski J, Sinha D, Jiang JC, Jazwinski SM, Hannun YA, Ogretmen B. Role of human longevity assurance gene 1 and C18-ceramide in chemotherapy-induced cell death in human head and neck squamous cell carcinomas. Mol Cancer Ther 6(2): 712–722, 2007. PMID:17308067

24. Brachtendorf S, Wanger RA, Birod K, Thomas D, Trautmann S, Wegner MS, Fuhrmann DC, Brüne B, Geisslinger G, Grösch S. Chemosensitivity of human colon cancer cells is influenced by a p53-dependent enhancement of ceramide synthase 5 and induction of autophagy. Biochim Biophys Acta Mol Cell Biol Lipids. 1863(10): 1214–1227, 2018. PMID: 30059758.

25. Kramer R, Bielawski J, Kistner-Griffin E, Othman A, Aleuc I, Ernst D, Kornhauser D, Hornemann T, Spassieva S. Neurotoxic 1-deoxysphingolipids and paclitaxel-induced peripheral neuropathy. FASEB J. 29(11): 4461–4472, 2015. PMID: 26198449

26. Prinetti A, Millimaggi D, D’Ascenzo S, Clarkson M, Bettiga A, Chigorno V, Sonnino S, Pavan A, Dolo V. Lack of ceramide generation and altered sphingolipid composition are associated with drug resistance in human ovarian carcinoma cells. Biochem J. 395(20: 311-318, 2006. PMCID: PMC1422777

27. Sänger N, Ruckhäberle E, Györffy B, Engels K, Heinrich T, Fehm T, Graf A, Holtrich U, Becker S, Karn T. Acid ceramidase is associated with an improved prognosis in both DCIS and invasive breast cancer. Mol Oncol. 9(1): 58–67, 2015. PMCID: PMC5528695

28. van Kruining D, Luo Q, van Echten-Deckert G, Mielke MM, Bowman A, Ellis S, Oliveira TG, Martinez-Martinez P. Sphingolipids as prognostic biomarkers of neurodegeneration, neuroinflammation, and psychiatric diseases and their emerging role in lipidomic investigation methods. Adv Drug Deliv Rev. 159: 232–244, 2020. PMCID: PMC7665829

29. Montfort A, Bertrand F, Rochotte J, Gilhodes J, Filleron T, Milhès J, Dufau C, Imbert C, Riond J, Tosolini M, Clarke CJ, Dufour F, Constantinescu AA, Junior NF, Garcia V, Record M, Cordelier P, Brousset P, Rochaix P, Silvente-Poirot S, Therville N, Andrieu-Abadie N, Levade T, Hannun YA, Benoist H, Meyer N, Micheau O, Colacios C, Ségui B. Neutral Sphingomyelinase 2 Heightens Anti-Melanoma Immune Responses and Anti-PD-1 Therapy Efficacy. Cancer Immunol Res. 9(5): 568–582, 2021. PMCID: PMC9631340.

30. Blaho VA, Galvani S, Engelbrecht E, Liu C, Swendeman SL, Kono M, Proia RL, Steinman L, Han MH, Hla T. HDL-bound sphingosine-1-phosphate restrains lymphopoiesis and neuroinflammation. Nature 523(7560): 342-346, 2015. PMCID: PMC4506268.

31. Galvani S, Sanson M, Blaho VA, Swendeman SL, Obinata H, Conger H, Dahlbäck B, Kono M, Proia RL, Smith JD, Hla T. HDL-bound sphingosine 1-phosphate acts as a biased agonist for the endothelial cell receptor S1P1 to limit vascular inflammation. Sci. Signal. 8(389): 79, 2015. PMID: PMC4768813

32. Zelnik ID, Kim JL, Futerman AH. The Complex Tail of Circulating Sphingolipids in Atherosclerosis and Cardiovascular Disease. J Lipid Atheroscler. 10(3): 268–281, 2021. PMCID: PMC8473959

33. Drobnik W, Liebisch G, Audebert FX, Frohlich D, Gluck T, Vogel P, Rothe G, Schmitz G. Plasma ceramide and lysophosphatidylcholine inversely correlate with mortality in sepsis patients. J Lipid Res. 44(4): 754–761, 2003. PMID: 12562829

34. Summers SA, Chaurasia B, Holland WL. Metabolic messengers: ceramides. Nat Metab 1:1051–1058, 2019. PMCID: PMC7549391

35. Zhao T, Li J, Wang Y, Guo X, Sun Y. Integrative metabolome and lipidome analyses of plasma in neovascular macular degeneration. Heliyon 9(10): e20329, 2023. PMCID: PMC10539639

36. Janneh AH, Kassir MF, Dwyer CJ, Chakraborty P, Pierce JS, Flume PA, Li H, Nadig SN, Mehrotra S, Ogretmen B. Alterations of lipid metabolism provide serologic biomarkers for the detection of asymptomatic versus symptomatic COVID-19 patients. Sci. Rep. 11(1): 14232, 2021. PMCID: PMC8270895.

37. Torretta E, Garziano M, Poliseno M, Capitanio D, Biasin M, Santantonio TA, Clerici M, Lo Caputo S, Trabattoni D, Gelfi C. Severity of COVID-19 Patients Predicted by Serum Sphingolipids Signature. Int J Mol Sci. 22(19): 10198, 2021. PMCID: PMC8508132.

38. Peterson LR, Xanthakis V, Duncan MS, Gross S, Friedrich N, Völzke H, Felix SB, Jiang H, Sidhu R, Nauck M, Jiang X, Ory DS, Dörr M, Vasan RS, Schaffer JE. Ceramide Remodeling and Risk of Cardiovascular Events and Mortality. J. Am. Heart Assoc. 7(10): e007931, 2018. PMCID: PMC6015315.

39. Nwabuo CC, Duncan M, Xanthakis V, Peterson LR, Mitchell GF, McManus D, Cheng S, Vasan RS. Association of Circulating Ceramides With Cardiac Structure and Function in the Community: The Framingham Heart Study. J. Am. Heart Assoc. 8(19): e013050, 2019. PMCID: PMC6806035.

40. Vasile VC, Meeusen JW, Medina Inojosa JR, Donato LJ, Scott CG, Hyun MS, Vinciguerra M, Rodeheffer RR, Lopez-Jimenez F, Jaffe AS. Ceramide Scores Predict Cardiovascular Risk in the Community. Arterio Thromb Vasc Biol. 41(4): 1558–1569, 2021. PMCID: PMC8939859.

41. Churchill RA, Gochanour BR, Scott CG, Vasile VC, Rodeheffer RJ, Meeusen JW, Jaffe AS. Association of cardiac biomarkers with long-term cardiovascular events in a community cohort. Biomarkers 29(4): 161–70, 2024. PMID: 38666319.

42. Poss AM, Maschek JA, Cox JE, Hauner BJ, Hopkins PN, Hunt SC, Holland WL, Summers SA, Playdon MC. Machine learning reveals serum sphingolipids as cholesterol-independent biomarkers of coronary artery disease. J Clin Invest. 130(3): 1363–1376, 2020. PMCID: PMC7269567.

43. Bhagat AA, Kalogeropoulos AP, Baer L, Lacey M, Kort S, Skopicki HA, Butler J, Bloom MW. Biomarkers and Strain Echocardiography for the Detection of Subclinical Cardiotoxicity in Breast Cancer Patients Receiving Anthracyclines. J Pers. Med. 13(12): 1710, 2023. PMCID: PMC10744645.

44. Jenkins RW, Clarke CJ, Lucas J, Shabir M, Wu BX, Simbari F, Mueller J, Hannun Yam Lazarchick J, Shirai K. Elevations of the role of Secretory Sphingomyelinase and Bioactive Sphingolipids as biomarkers in Hemophagocytic Lymphohistiocytosis. Am. J. Hematology 88(11): E265-72, 2013. PMCID: PMC4348111.

45. Hammad SM, Pierce JS, Soodavar F, Smith KJ, Al Gadban MM, Rembiesa B, Klein RL, Hannun YA, Bielawski J, Bielawska A. Blood sphingolipidomics in healthy humans: impact of sample collection methodology. J Lipid Res. 51(10): 3074–3087, 2010. PMCID: PMC2936747

46. Luu AZ, Chowdhury B, Al-Omran M, Teoh H, Hess DA, Verma S. Role of Endothelium in Doxorubicin-Induced Cardiomyopathy. J ACC Basic Transl Sci 3(6): 861–870, 2018. PMCID: PMC6314956

47. Murata T, Yamawaki H, Yoshimoto R, Hori M, Sato K, Ozaki H, Karaki H. Chronic effect of doxorubicin on vascular endothelium assessed by organ culture study. Life Sci. 69(22): 2685–2695, 2001. PMID: 11712671

48. Lee VW, Harris DC. Adriamycin nephropathy: a model of focal segmental glomerulosclerosis. Nephrology. 16: 30–38, 2011. PMID: 21175974

49. Damodar G, Smitha T, Gopinath S, Vijayakumar S, Rao Y. An evaluation of hepatotoxicity in breast cancer patients receiving injection Doxorubicin. Ann Med Health Sci Res. 4(1): 74–9, 2014. PMID: 24669335

50. Cardinale D, Colombo A, Bacchiani G, Tedeschi I, Meroni CA, Veglia F, Civelli M, Lamantia G, Colombo N, Curigliano G, Fiorentini C, Cipolla CM. Early detection of anthracycline cardiotoxicity and improvement with heart failure therapy. Circulation 131(22): 1981–1988, 2015. PMID: 25948538.

51. Cardinale D, Colombo A, Lamantia G, Colombo N, Civelli M, De Giacomi G, Rubino M, Veglia F, Fiorentini C, Cipolla CM. Anthracycline-induced cardiomyopathy: clinical relevance and response to pharmacologic therapy. J Am Coll Cardiol. 55(3): 213–220, 2010. PMID: 20117401

